# A PATHWAY FOR T3 SIGNALING IN THE BRAIN TO IMPROVE THE VARIABLE EFFECTIVENESS OF THERAPY WITH L-T4

**DOI:** 10.1101/2022.08.17.504300

**Authors:** Federico Salas-Lucia, Csaba Fekete, Richárd Sinkó, Péter Egri, Kristóf Rada, Yvette Ruska, Barbara Bocco, Tatiana Fonseca, Balázs Gereben, Antonio C. Bianco

**Author notes:** Both authors contributed equally to the work (Federico Salas-Lucia, Csaba Fekete). **Corresponding authors:** Antonio C. Bianco, MD, PhD - Section of Endocrinology, Diabetes & Metabolism, University of Chicago Medical Center, 5841 S. Maryland Ave. MC1027, Room M267 | Chicago, IL 60637, Balázs Gereben, DVM, PhD – Laboratory of Molecular Cell Metabolism, Institute of Experimental Medicine, 43 Szigony str., Budapest, 1083 Hungary.

## Abstract

The effectiveness of therapy for hypothyroidism with levothyroxine (L-T4) depends on patients’ ability to activate T4 to T3 —altered in carriers of a common deiodinase polymorphism (Thr92Ala-DIO2). Some patients that exhibit impaired mood and cognition improve with liothyronine (L-T3), but the underlying mechanisms remain unknown. Here we show that the T3-indicator mouse carrying the Thr92Ala-DIO2 polymorphism exhibits a hippocampal-specific reduction in T3 activation and signaling that limits the effectiveness of L-T4 therapy. To understand the L-T3 effect, we used a compartmentalized microfluid device and identified a novel neuronal pathway of T3 transport and action that involves axonal T3 uptake into clathrin-dependent, endosomal/non-degradative lysosomes (NDLs). NDLs-containing T3 are retrogradely transported via microtubules, delivering relatively large amounts of T3 to the cell nucleus, doubling the expression of the T3-responsive reporter gene. The NDLs also contain the monocarboxylate transporter 8 (Mct8) and the type 3 deiodinase (Dio3), which transports and inactivates T3, respectively. Notwithstanding, T3 gets away from degradation because D3 active center is in the cytosol. These findings provide (i) a basis for the variable effectiveness of L-T4 therapy, (ii) a pathway for L-T3 to reach neurons, and (iii) resolve the paradox of T3 signaling in the brain amid high D3 activity.

## Introduction

The standard of care for the treatment of hypothyroidism is the daily administration of levothyroxine (L-T4) tablets. Symptoms are resolved for most patients once serum TSH levels have returned to the normal range. Nonetheless, even appropriately treated patients with hypothyroidism might not fully recover and remain symptomatic with detriments to cognition, mood, and quality of life (*1*). Studies in Europe and the US point to up to 15% of the patients with hypothyroidism experiencing residual cognitive symptoms, many times referred to as “brain fog” by patients. While non-thyroid-related reasons might contribute to these residual symptoms, incomplete normalization of thyroid hormone (TH) homeostasis and TH action in the brain also plays a critical role in this process (*2*).

TH are crucial for brain development and influence brain function throughout life (*3–5*). However, the complex architectural organization of the brain and unique properties of each cell type pose a challenge to the complete understanding of the mechanisms that regulate local TH signaling (*6*). Neurons are an important TH target in the brain as they express the highest levels of TH receptors (TRs) (*7*). The blood-brain barrier (BBB) and glial cells also play a role in TH signaling in the brain, regulating the amount of TH that reaches neurons through selective transport and metabolism (*8*, *9*). As a result, the human brain responds promptly to minor fluctuations in TH signaling by changing the expression of T3-responsive genes (*10*). TH transporters seem to play a critical role in T3 signaling as showcased by the profound brain hypothyroidism observed in boys carrying mutations in the monocarboxylate transporter 8 (MCT8, SlC16A2). The resulting Allen Herndon Dudley syndrome is marked by severe and irreversible neurological damage (*11*) due to reduced TH availability to neurons.

Plasma T3 can reach neurons via cellular transporters located in the BBB (*12*, *13*), including MCT8—other species-specific transporters also play a role—but most T3 bound to TRs in the brain is originated from local D2 (*7*, *14*), the enzyme that catalyzed conversion of T4 to T3. Within the brain, D2 is expressed in glial cells, astrocytes and tanycytes, but not in neurons (*15*, *16*). Astrocytes are intimately related to neurons (hundreds of thousands of neuronal synapses per astrocyte); an array of metabolites (and even mitochondria) are known to be preferentially exchanged between astrocytes and neurons (*17*). Hence, the concept that the brain responsiveness to L-T4 is mediated by astrocyte-derived T3 production and transfer to neighboring neurons is well accepted (*9*, *18*). Accordingly, a mouse with glial-cell selective inactivation of the gene encoding D2 (Dio2) exhibits a mood and cognitive phenotype typical of hypothyroidism (*19*).

A common polymorphism in the DIO2 gene (Thr92Ala-DIO2), which alters the intracellular localization of the enzyme and reduces D2 activity by ~20%, has been associated with a worse response to L-T4 in the treatment of hypothyroidism, and clinical improvement when liothyronine (L-T3) was added to L-T4 therapy (*20–22*). Although not universally observed (*23*), these findings highlight the potential for the D2 pathway to affect normal TH signaling in the brain and modify the effectiveness of therapy with L-T4. Indeed, a mouse carrying the Thr92Ala-Dio2 polymorphism exhibits a transcriptome and a cognitive phenotype suggestive of a localized reduction in TH signaling in the brain—the mice also responded positively to therapy with L-T3 (*24*).

It is unexpected that treatment with L-T3 can be effective given that neurons express high levels of the TH inactivating type 3 deiodinase (Dio3) (*25*). Based on all we know about D3, its role in neurons should function as a barrier to incoming T3, minimizing or preventing T3 signaling (*26*). And yet, we know that brain TRs are nearly fully occupied with T3 (*7*). Thus, it is not clear how T3 escapes from D3-mediated degradation in neurons and reaches the nucleus of these cells, and consequently how treatment with small amounts of L-T3 can be effective in restoring brain-localized hypothyroidism in mice and humans.

Here we studied the effectiveness of treatment with L-T4 in restoring TH signaling in different brain areas of a mouse carrying the Ala92-Dio2 allele. We also defined the cellular mechanisms underlying T3 actions in neurons, with broad implications for L-T3 based therapies for hypothyroidism.

## Results

### Hippocampal Responsiveness to L-T4 is Impaired in the Ala92-Dio2 Mouse

Thr92- and Ala92-Dio2 mice were crossed with the Thyroid Hormone Action Indicator (THAI) indicator mouse (*27*)—which ubiquitously expresses a luciferase (Luc) reporter gene driven by three in-tandem TH response elements (TRE)—to produce homozygous Thr92- and Ala92-Dio2 mice expressing the THAI reporter gene. Whereas Luc-mRNA levels (the read-out for TH action) in different parts of the brain were similar between Thr92- and Ala92-Dio2 mice, in the cerebellum Luc-mRNA was lower in the Ala92-Dio2 mice (**Table 1**). Next, these mice were made hypothyroid (low iodine diet combined with anti-thyroid drugs) and some, as indicated, were treated with daily doses of L-T4 (1.7 or 1.9 μg/100g BW/day via gavage) (**Fig. 1A**). Hypothyroid mice had undetectable serum T4 and T3 and elevated serum TSH levels, regardless of their genotype (**Fig. 1B-D**). Treatment with 1.7 μg L-T4/100g BW/day elevated serum T4 levels above controls while the serum T3 levels remained slightly below control values; serum TSH levels were normalized (**Fig. 1B**). Treatment with the higher dose of L-T4 slightly elevated serum T3 levels even further and lowered serum TSH levels, while maintaining elevated serum T4 levels (**Fig. 1B-D**). This pattern of thyroid function tests (i.e. normal serum TSH with slightly elevated serum T4 levels and low/normal serum T3 levels) is frequently observed in LT4-treated patients with hypothyroidism (*28*).

**Fig. 1.**
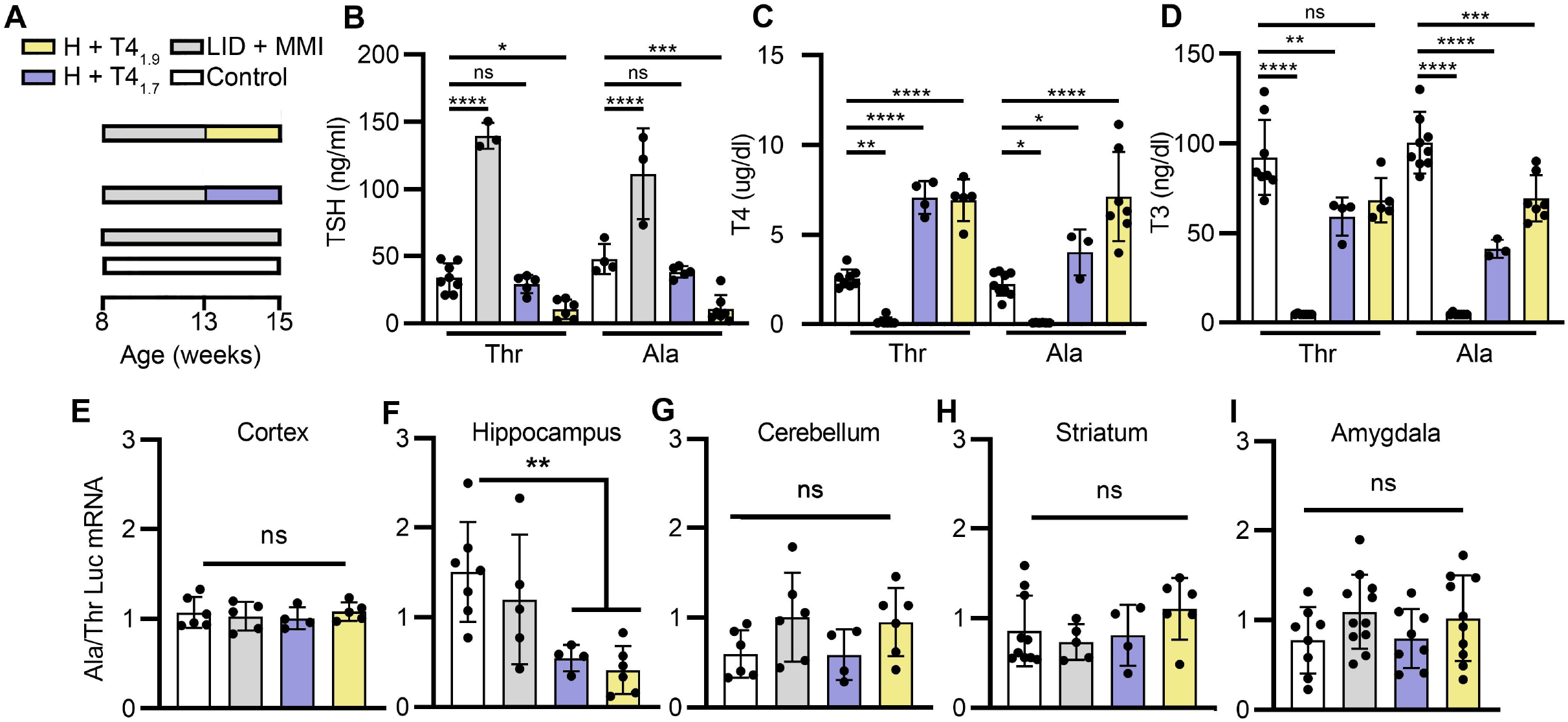
Hippocampal responsiveness to L-T4 is impaired in the Ala92-Dio2 mice. **A.** Diagram shows the experimental groups studied; colored horizontal bars show different treatments. **B-D**. Plasma TSH (**B**), T4 (**C**), and T3 (**D**) in mice undergoing the indicated treatment. **E-I**. Ala92-Dio2/Thr92-Dio2 ratios of the Luc mRNA levels in the indicated areas and treatments. Note how in the hippocampus, the Ala92-Dio2 mice expressed significantly lower Luc mRNA levels. Values are mean ± SD of 3-10 independent experiments; \**P* <0.05, \*\**P* < 0.01, \*\*\**P* < 0.001; ns: non-significant.

**Table 1:** Summary of the changes in Luc mRNA expression in the Thr92-Dio2-THAI and Ala92-Dio2-THAI mice undergoing the indicated treatments.

| Brain area | Basal euthyroid | Hypothyroid |  | Hypothyroid + T4<sub>1.70</sub> |  | Hypothyroid + T4<sub>1.9</sub> |  |
| --- | --- | --- | --- | --- | --- | --- | --- |
|  | Ala92 | Thr92 | Ala92 | Thr92 | Ala92 | Thr92 | Ala92 |
| Cortex | ↔ | ↓*** | ↓** | ↔ | ↔ | ↑** | ↑** |
| Striatum | ↔ | ↓**** | ↓*** | ↔ | ↔ | ↔ | ↔ |
| Cerebellum | ↔ | ↓*** | ↓* | ↓* | ↓* | ↓* | ↔ |
| Hippocampus | ↔ | ↓* | ↓* | ↔ | ↓* | ↑* | ↓* |
| Amygdala | ↔ | ↔ | ↔ | ↔ | ↔ | ↔ | ↔ |

| Brain area | Basal euthyroid | Hypothyroid |  | Hypothyroid + T4<sub>1.70</sub> |  | Hypothyroid + T4<sub>1.85</sub> |  |
| --- | --- | --- | --- | --- | --- | --- | --- |
|  | Ala92 | Thr92 | Ala92 | Thr92 | Ala92 | Thr92 | Ala92 |
| Cortex | ↔ | ↓*** | ↓** | ↔ | ↔ | ↑** | ↑** |
| Striatum | ↔ | ↓**** | ↓*** | ↔ | ↔ | ↔ | ↔ |
| Cerebellum | ↔ | ↓*** | ↓* | ↓* | ↓* | ↓* | ↔ |
| Hippocampus | ↔ | ↓* | ↓* | ↔ | ↓* | ↑* | ↓* |
| Amygdala | ↔ | ↔ | ↔ | ↔ | ↔ | ↔ | ↔ |

The respective Luc-mRNA changes in the cerebral cortex, striatum, cerebellum, hippocampus, and amygdala of the Thr92- and Ala92-Dio2 mice are shown in **Table 1**. Hypothyroidism caused a reduction in the Luc-mRNA levels in the cerebral cortex (~30%), cerebellum (~60%), hippocampus (~40%), and striatum (80%), but not in the amygdala (**Table 1**). Treatment with the lower dose of L-T4 restored the euthyroid Luc-mRNA pattern in most areas of the Thr92- and Ala92-Dio2, but not in the Thr92- and Ala-92-Dio2 cerebellum and in the Ala92-Dio2 hippocampus (**Table 1**). The higher L-T4 dose increased Luc-mRNA levels above controls in the Thr92- and Ala92-Dio2 cerebral cortex and in Thr92-Dio2 hippocampus, whereas in the striatum the Luc-mRNA levels remained similar to controls. At the same time, Luc-mRNA levels remained lower in the Ala92-Dio2 hippocampus (**Table 1**). The Ala92/Thr92-Dio2 Luc-mRNA ratios illustrate the differences between the changes in the Thr92- and Ala92-Dio2 mice (**Fig. 1E-I**).

Whereas the cerebral cortex, cerebellum, striatum, and amygdala Luc-mRNA behaved similarly, the Ala92-Dio2 hippocampus Luc-mRNA failed to fully respond to treatment with L-T4 (**Fig. 1F**).

### Ala92-Dio2 Astrocytes Have Limited Ability to Activate T4 to T3

Ala92-D2 is slower at producing T3 in proliferating myoblasts and pituitary cells transiently expressing Ala92-Dio2 (*29*), and exhibits about 20% less enzymatic activity in HEK-293 cells stably expressing Ala92-Dio2 (*30*). Here we isolated and established primary cultures of astrocytes obtained from Thr92- and Ala92-Dio2 mice, which allowed us to study Ala92-D2 at endogenous brain levels. The astrocytes were obtained from the cerebral cortex, cerebellum, and hippocampus (**Fig. 2A**), and after about 10 days in culture were incubated with T4^I125^ for 24h for subsequent measurement of T3^I125^ production. Astrocytes obtained from the Ala92-Dio2 cerebral cortex and cerebellum produced similar amounts of T3^I125^, but the hippocampal astrocytes produced ~50% less T3^I125^ (**Fig. 2B**), which explains the findings of reduced Luc-mRNA in the hippocampus of L-T4-treated Ala92-Dio2 mice.

**Fig. 2.**
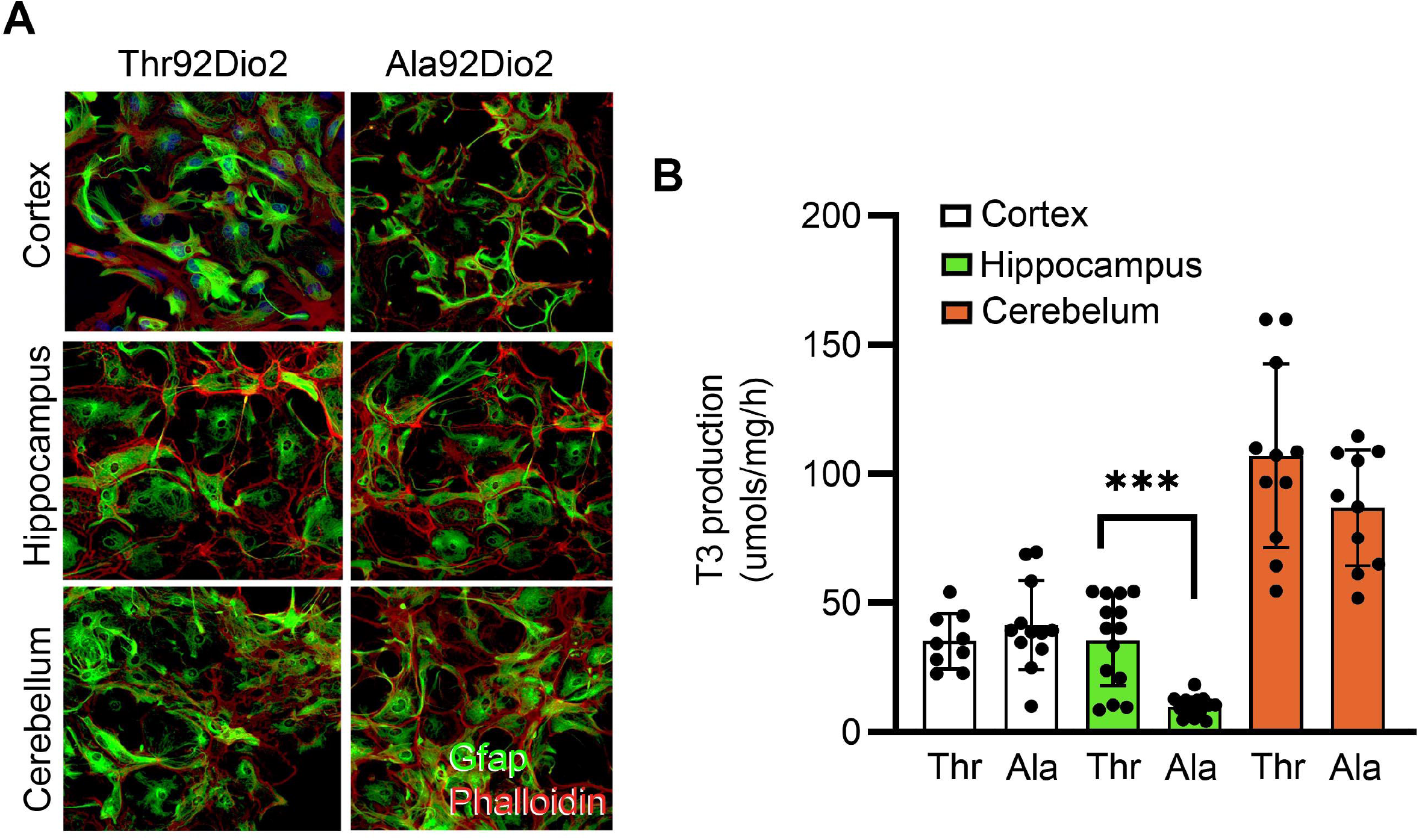
Hippocampal Ala92-Dio2 astrocytes have impaired ability to activate T4 to T3. **A.** Collage showing primary cultured astrocytes stained for Gfap (green) and phalloidin (red), obtained from the indicated brain areas of the Thr92- and Ala92Dio2 mice. **B.** Thr92- and Ala92 T3 production in primary astrocytic cultures obtained from the indicated areas.

### Neuronal Response to T3 Involves MCT8 and Retrograde TH Transport

Short-term injections of L-T3 fully normalizes the cognitive phenotype of the Ala92-Dio2 mouse, indicating that the impaired T4 to T3 conversion can be corrected with exogenous LT3 (*30*). In fact, the brain responds rapidly—within hours—to injections of L-T3 (*31*), including the Luc-mRNA levels in the THAI mouse (**Fig. S4A,B**), and promptly restores TH signaling in LT4-treated rats with hypothyroidism (*32*). However, the underlying mechanism of such a response remains elusive. How is it that sufficient *plasma* T3 can rapidly exit the capillaries, diffuse and enter neurons to reach the nuclear TRs amid very high levels of D3 expressed in neurons? D3 rapidly inactivates T3 to T2 in neurons (*33*), and thus it is likely to jeopardize the clinical effectiveness of any therapy for hypothyroidism that contains L-T3.

To obtain a mechanistic basis for such a scenario, we next deconstructed the brain’s architecture using a compartmentalized microfluidic chamber (MC), which purposely keeps the neuronal cell bodies and their long distal axons in two physically separated chambers. Primary cortical neurons (PCN) obtained from E16.5 mouse embryos were seeded in the cell side (MC-CS) and cultured for up to 13 days (**Fig. S1A**). During this period, distal axons grew from parental cell bodies into the axonal side of the microchamber (MC-AS) (**Fig. S1B**). By day *in vitro* (DIV) ~7, PCN had crossed the 450 μm long microgroove channels and by DIV ~10 densely populated the MC-AS (**Fig. S1B; Fig. 3A**). The addition of calcein fluorescence to the MC-CS revealed labeled axons (1-3 axons/channels) reaching the MC-AS (inset in **Fig. S1C**).

**Fig. 3.**
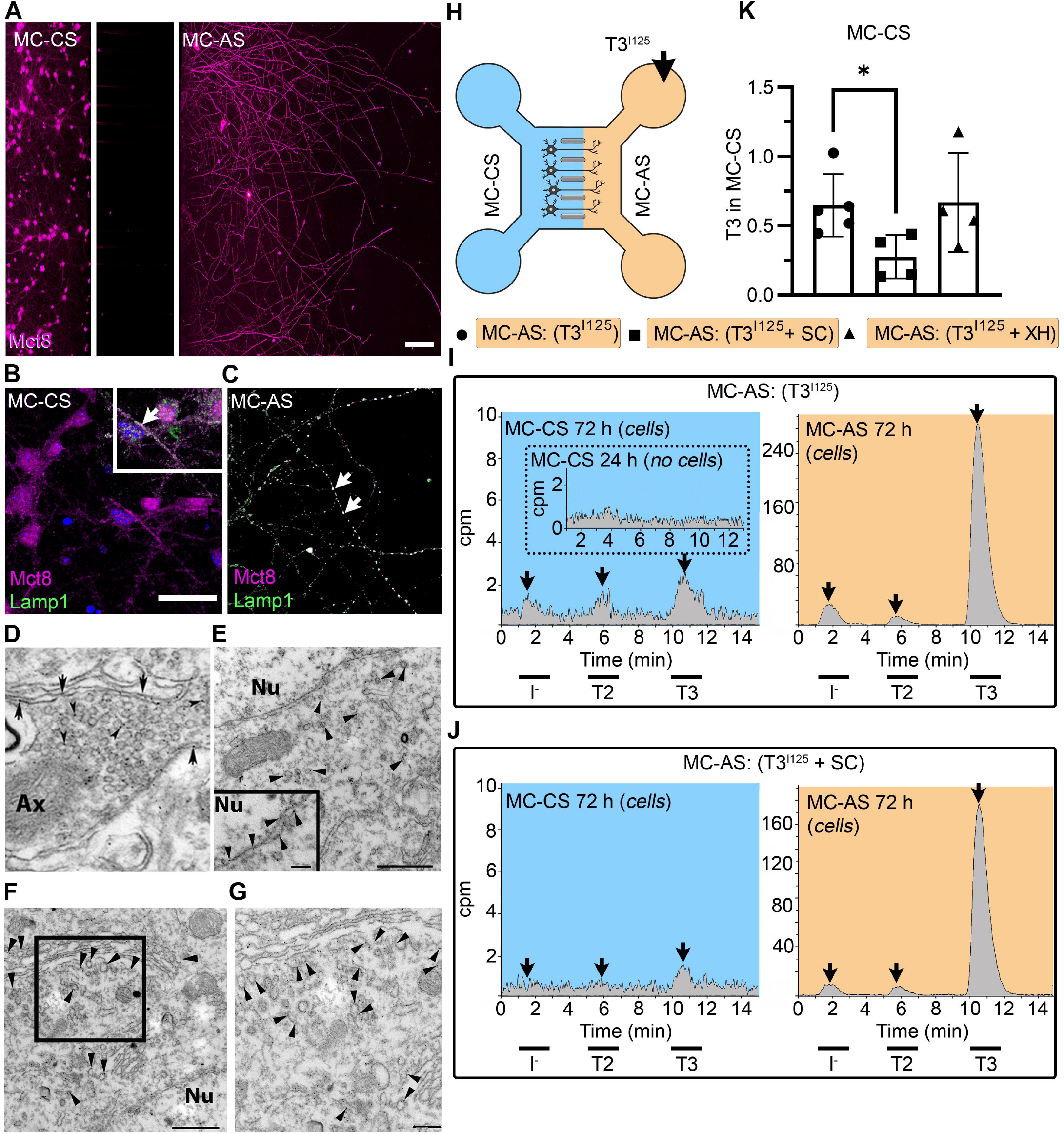
T3 is taken up and retrogradely transported through neuronal long distal axons. **A-C.** Immunofluorescence of cortical neurons in the compartmentalized device using the indicated antibodies **A.** Collage of a compartmentalized culture stained for MCT8 (magenta). **B-C.** MCT8 staining in magenta, Lamp1 in green; colocalization is indicated as white. **B** In the MC-CS and **C** in the MC-AS. **D.** MCT8-immunoreactivity (silver grains) was present in the outer cell membrane of neuronal elements, and in the membrane of vesicles of axonal profiles (arrows and arrowheads, respectively), on the nuclear membrane (inset in **E**), in vesicles close to the nucleus of the neurons (**E**), and the trans and cis Golgi apparatus (**F**). **G.** Detail of the squared area in **F**. **H**. T3^I125^ was applied in the MC-AS. **I**. After 72 h, T3^I125^ was detected in the MC-CS medium (no T3^I125^ was detected after 24h; inset in the blue panel). **J**. Effect of 2μM SC on T3 uptake and transport. The size of the T3^I125^ peak in the MC-CS decreased after 72 h compared to **I**. **K**. Quantitation of T3^I125^ transported to the MC-CS medium under the indicated conditions. The Y-axis in % T3^I125^ in the MC-CS medium vs. T3^I125^ added to the MC-AS medium. Values are mean ± SD of 4 – 5 independent experiments; \**P* < 0.05 in comparison with T3^I125^ incubation. Abbreviations: Ax, axon; Nu, nucleus. Scale bars are 25 μM on A-E and G, 150 μM on F, 500 nm on J, K, and M, and 200 nm on L.

Immunofluorescent studies showed that by DIV ~10, PCNs express MCT8 in both cellular compartments (**Fig. 3A-C**). Immunostaining for MCT8 revealed that MCT8 is present in the cell bodies and short processes and the long-distant axons (**Fig. 3B,C**). MCT8 colocalizes with Lamp1, a marker for endosome- and non-degradative lysosome (NDL)-like organelles (*34*), in cell bodies and axons (inset in **Fig. 3B,C**). MCT8 distribution to the axonal vesicles and plasma membrane was also detected through immune-electron microscopy (**Fig. 3D**). Furthermore, MCT8 was detected in the nuclear membrane, and the trans and cis Golgi apparatus, (**Fig. 3E-G**).

The addition of T3^I125^ to the MC-AS (**Fig. 3H**) led to T3 uptake and transport through neuronal long distal axons. Whereas only background radioactivity was detected in the medium in the MC-CS (inset in **Fig. 3I**, blue panel) after 24h, later (after 72h) about 0.5-1.0 % of T3^I125^ was detected in the MC-CS (**Fig. 3I**, blue panel), illustrating that retrograde transport occurred, i.e., MC-AS→MC-CS. Small amounts of T2^I125^ and I^125^ were also present, likely the result of T3^I125^ metabolism. Given the concentration of T3 in the medium (2.6 nM) we estimate that between 0.8 and 1.6 fmols of T3 were retrogradely transported during the 72h incubation by the approximately 300 neurons that crossed the MC-CS into the MC-AS.

To test whether the retrograde T3^I125^ transport was mediated via MCT8, we next added 2 μM Sylicrhistin (SC; a selective MCT8 inhibitor) (*35*) to the MC-AS and saw that it decreased the transport of T3^I125^ (**Fig. 3K**) as evidenced by the smaller size of the T3^I125^ peaks detected in the MC-CS (**Fig. 3J**, blue panel). We also used the same setup to test whether retrograde T3^I125^ transport occurred in other types of neurons (**Fig. S2A,B**). We isolated neurons from postnatal day (P) 2 rat dorsal root ganglia (DRG), which also express MCT8 (**Fig. S2D**). Similar to cortical neurons, DIV ~10 DRG cells exhibited MC-AS→MC-CS transport of T3^I125^, inhibited by 2 μM SC (**Fig. S2C**). Altogether, these results show that T3 is retrogradely transported through axons and released on the opposite cellular compartment (MC-AS → MC-CS), with the involvement of MCT8.

### Retrograde T3 Transport Via Neuronal Endosomes/NDL

It is well known that a retrograde transport system exists in neurons based on neuronal endosomes/NDL (*36*). Thus, we hypothesized that the transport of T3 uses this shuttle mechanism. To test if this was the case, we isolated and cultured DIV ~5 PCNs, which were then loaded with T3^I125^ for 24 h. Cells were then harvested and processed through iodixanol gradient ultracentrifugation for isolation of subcellular fractions containing endosomes/NDL, including fraction one (F1) that was enriched with the endosomes/NDL (**Fig. 4A**). The resulting fractions were then resolved through UPLC and it was clear that most T3^I125^ was contained in F1, with some spillover to F2 (**Fig. 4B**).

**Fig. 4.**
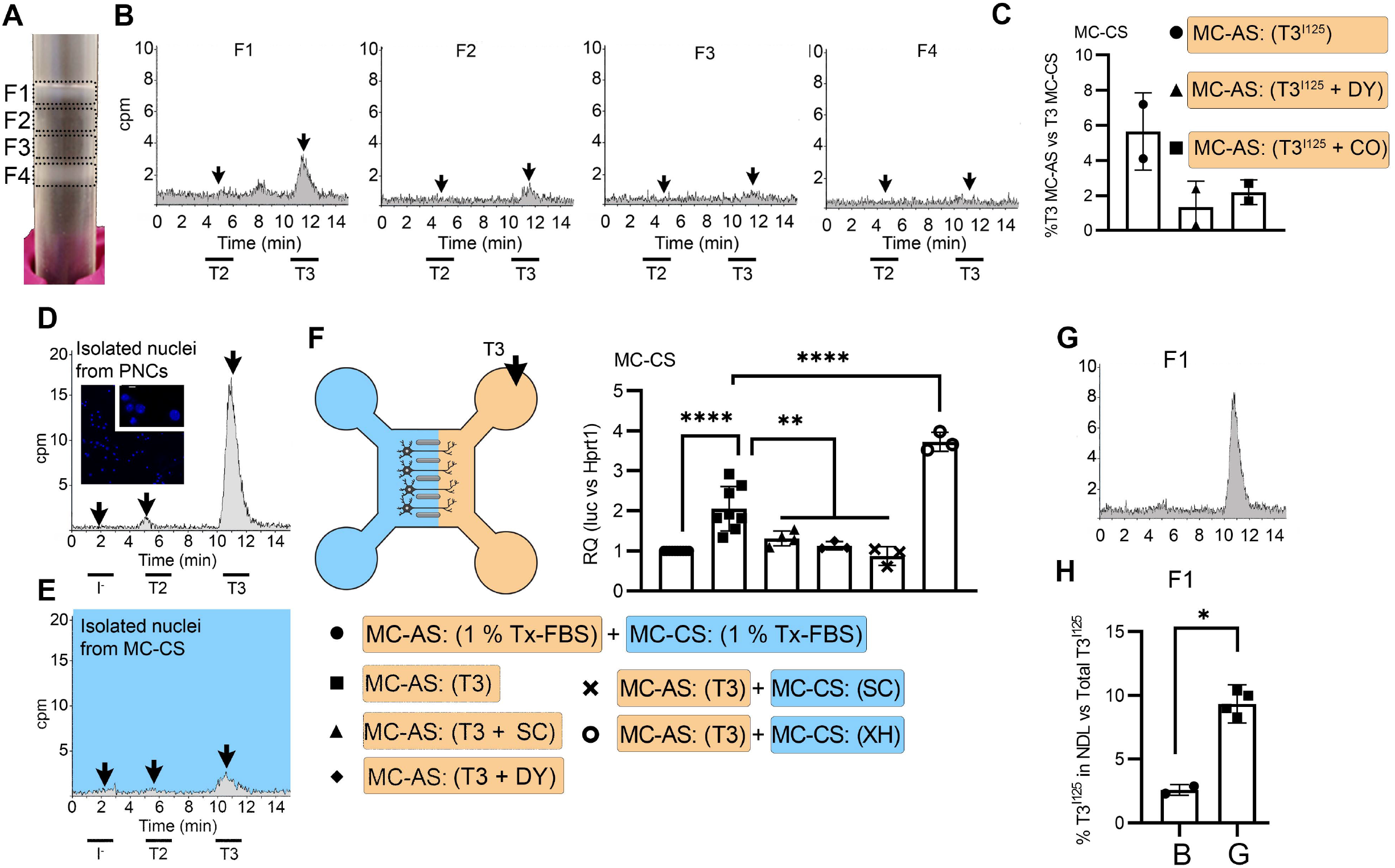
T3 is in neuronal endosomes/NDL; retrograde transport depends on clathrin-mediated endocytosis and microtubules and initiates TH signaling. **A**. Gradient column after ultracentrifugation, the resulting four fractions are indicated. **B.** Chromatograms of the medium after the PNCs were incubated with T3^I125^ for 24 h, and of the four fractions after ultracentrifugation. **C.** Quantitation of T3^I125^ retrogradely transported into the MC-CS medium under the indicated conditions. The Y-axis in % T3^I125^ in the MC-CS vs. T3^I125^ added to the MC-AS. **D.** ^~^ 9 x 10^6^ PNCs were loaded with T3^I125^ for 24h and processed for nuclei isolation, which were subsequently studied through UPLC, showing an accumulation of T3^I125^. **E**. Same as in **D**, except that the T3^I125^ was applied in the MC-CS and the nuclei isolated from ^~^200.000 neurons from the MC-AS (pool of 10 microchambers). **F.** 10 nM T3 was applied in the MC-AS for 24 h. Bar graph shows the quantitation of the Luc mRNA levels of the cells in the MC-CS under the indicated conditions. **G**. Same as in **B**, except that endosomes/NDL were isolated in the presence of 2 μM SC. **H.** Quantitation of the T3^I125^ from the chromatograms found in fraction 1 in **B** and **G**. The Y-axis in % T3^I125^ in the F1 vs. T3^I125^ was added to the medium. Values are mean ± SD of 3 – 8 independent experiments; \**P* < 0.05, in comparison with 10 nM T3 incubation, \*\**P* < 0.01, \*\*\**P* < 0.001 in comparison with 1 % Tx FBS incubation.

Endosomes/NDL can be formed through clathrin-mediated endocytosis and be actively transported throughout microtubules (*37*). To find out whether these elements were involved in the formation of endosomes/NDL containing T3, we co-incubated the MC-AS with T3^I125^ and either 20 μM Colchicine (CO, an inhibitor of microtubule formation)or 80 μM dynasore (DY), an inhibitor of clathrin-mediated endocytosis (*38*). The use of CO or DY reduced the amount of T3^I125^ detected in the MC-CS **(Fig. 4C**). Similarly, when used with DRGs neurons, CO reduced the amount of radiation detected in the MC-CS (**Fig. S2C**).

### Retrograde T3 transport initiates TH signaling in cerebral cortex neurons

It is logical to assume that some of the retrogradely transported T3 ends up in the nucleus of the neurons where it can initiate TH signaling. We first looked for T3^I125^ in the nuclei of approximately 400,000 neurons after T3^I125^ was added to MC-AS. These neurons include the approximately 300 neurons that crossed the bridge between MC-AS and MC-CS and mediate the retrograde transport of T3^I125^. Indeed, already after 24h a clear peak of T3^I125^ could be identified in the nuclear fraction of these neurons (**Fig. 4E**). The identity of the T3^I125^ peak was confirmed because it comigrated with a much more prominent T3^I125^ peak obtained from nuclei of 8-9×10^6^ cells directly labeled with T3^I125^ (**Fig. 4D**). We next utilized the T3 concentration in the medium (2.7nM) and the nuclei/medium ratio of T3^I125^ (~0.0015) and estimated that the nuclei in the neurons contain approximately 0.75 ng T3/mg DNA that originated from the retrograde transport from the MC-AS. This figure is similar to what was obtained in the rat’s cerebral cortex after injection of T3^I125^ (*7*), suggesting that the retrograde transport of T3 is of physiological relevance.

The presence of T3^I125^ in the nuclei of the neurons indicates that the retrograde T3 transport has the potential to affect TH signaling via the initiation of T3-dependent regulation of gene expression. To test if this was the case, we next isolated cortical neurons from THAI E16.5 embryos (*27*). MC-AS was incubated with 10 nM T3 (200 pM free T3), and 24 h later, neurons in the MC-CS were harvested and processed for Luc mRNA determination (**Fig. 4F**). The addition of 10 nM T3 to the MC-AS resulted in a 2.2 ± 0.5-fold increase in Luc mRNA levels (**Fig. 4F**). However, coincubation of T3 with 2 μM SC in the MC-AS significantly reduced T3 induction of Luc mRNA, highlighting the importance of MCT8 in this mechanism (**Fig. 4F**).

We also tested whether the addition of DY to the MC-AS interfered with the T3 induction of Luc. Indeed, after 24h of the addition of T3, induction of Luc mRNA was blunted in the presence of DY (**Fig. 4F**), confirming that clathrin-dependent endosomal T3 uptake is involved in T3 action in neurons. A corollary is that T3 is taken up in the MC-AS by endosomal/NDL, transported via microtubules to the MC-CS, and released to the cell nucleus where it regulates gene expression.

### MCT8 Modulates the Exit of T3 from the Endosomal/NDL

Not much is known about how contents in the endosomal/NDL are transferred to the cell nucleus, and here we studied whether MCT8 could have a role in this process. This was first tested *in vitro* by isolating T3^I125^-loaded endosomes/NDL in the presence or absence of 2 μM SC. The presence of SC during isolation was associated with 4-fold retention of T3^I125^ inside the F1 endosomal/NDL (**Fig. 4G,H**), suggesting that MCT8 mediates the release of T3 from the endosomes/NDL. Second, we tested whether the presence of SC in MC-CS could affect TH signaling initiated by the retrograde transport of T3 from the MC-AS. While the addition of 10 nM T3 in the MC-AS doubled Luc mRNA levels after 24h, the presence of 2 μM SC in the MC-CS blunted Luc mRNA induction by T3 (**Fig. 4F**), indicating that MCT8 plays a role in the release of T3 to the cell nucleus and initiation of TH signaling.

### T3 is not Metabolized in MC-AS Despite the Presence of D3 in Axons

Immunofluorescent studies showed that by DIV ~10, PCNs express D3 in both cellular compartments (**Fig. 5A,B**). To visualize D3, we used a D3-specific antibody directed against the molecule’s C-end (D3_250-300_). Staining with α-D3_250-300_ revealed that D3 is present in the cell bodies and short processes (**Fig. 5A**) and the long-distant axons (**Fig. 5B**), where the D3 signal displays a dotted pattern. This was reminiscent of our previous observations that D3 is present in early endosomes and constantly recycles with the plasma membrane (*33*, *39*, *40*). To test if D3 was compartmentalized in this system as well, we looked for colocalization of D3 with Lamp-1. We found that D3 co-localizes with Lamp1 in the cell bodies and axons (**Fig. 5A,B**). There was also a weak D3 signal in the cell nucleus (**Fig. 5A)**, which is in agreement with our previous studies showing that D3 sorts to the neuronal nucleus only during hypoxic conditions (*33*).

**Fig. 5.**
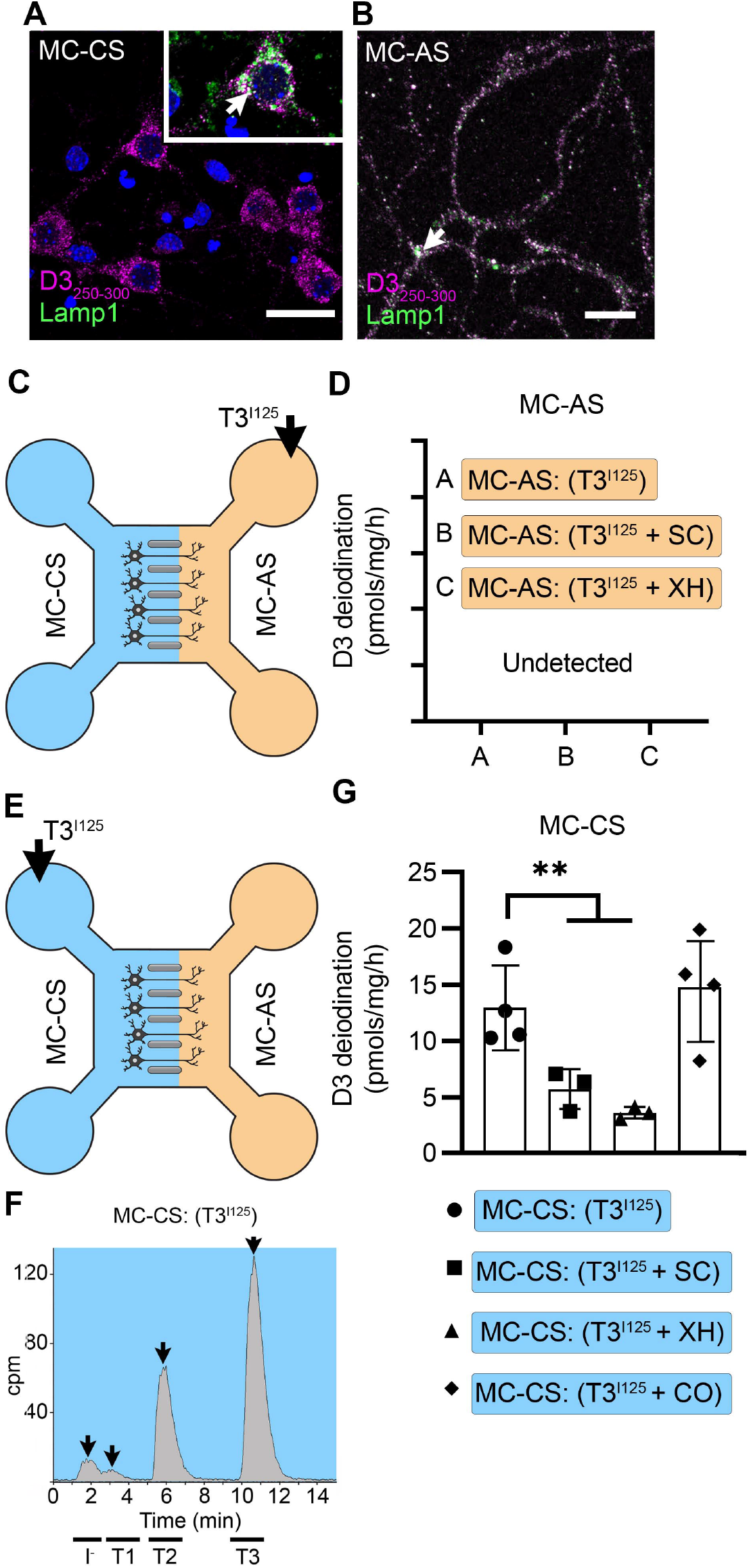
T3 is not metabolized by axonal D3. **A.** In the MC-CS D3 staining in magenta and Lamp1 staining in green; colocalization is indicated as white and shown in the inset (arrow). **B**. In the MC-AS, D3 staining in magenta and Lamp1 staining in green; colocalization is indicated as white (arrow). **C,D**. No D3-mediated T2^I125^ production was detected in the MC-AS. However, a high D3-mediated T2^I125^ production was detected in the MC-CS (**E-G**), which was reduced by exposure to 2 μM SC or 6 μM XH but not by exposure to 80 μM DY. Values are mean ± SD of 3 – 4 independent experiments; \*\**P* < 0.01 in comparison with T3^I125^ incubation.

Despite the abundant presence of D3 in the long-distant axonal network, the addition of T3^I125^ to MC-AS (**Fig. 5C**) for up to 72 h revealed no deiodination of T3^I125^; only background amounts of T2^I125^ were detected in the medium, equivalent to when no cells were added (**Fig. 5C,D**). Moreover, the addition of 6 μM of the D3 inhibitor XH to the MC-AS also did not affect the retrograde transport of T3^I125^ (**Fig. 5D**). D3 is a transmembrane protein with the active center of the enzyme located in the cytosol (*41*). Thus, it is conceivable that most T3 that is taken up by axons ends up in the endosomal/NDL vesicles rather than in the cytosol, where it would be easily metabolized by D3.

Notably, a different scenario altogether was identified in the body of the neurons. The addition of T3^I125^ to the MC-CS (**Fig. 5E**) resulted in a prominent peak of T2^I125^, which reflects the uptake of T3^I125^ to the cytosol, metabolism, and release of T2^I125^ to the medium (**Fig. 5F**). The rate of T3 metabolism was high, consuming approximately ¼ of the added T3^I125^ at every 24h. The addition of 2 μM SC or 6 μM XH to the MC-CS markedly reduced the metabolism of T3^I125^ (**Fig. 5G**), evidenced by the decrease in the peaks of T2^I125^ found in the MC-CS. As expected, the addition of SC or XH in the MC-AS did not affect the metabolism of T3^I125^ in the MC-CS (**Fig. 3A, B**), indicating that these drugs are not transported across compartments. These data suggest that T3 entering the neuronal cell body via MCT8 is rapidly targeted for inactivation via D3.

### D3 in the MC-CS Modulates TH Signaling Initiated by Retrograde Transport of T3

The fact that T3 in the endosomal/NDL vesicles is transferred to the cell nucleus to initiate TH signaling in the neighborhood of high D3 activity in the neuronal cell bodies raises the possibility that those D3 enzymes could modulate TH signaling. This was investigated by measuring TH signaling initiated by retrogradely transported T3 in the presence or absence of 6 μM XH in the MC-CS. Remarkably, XH enhanced the T3 induction of Luc mRNA levels by 1.8-fold (**Fig. 4F**). These results suggest that D3 activity in the neuronal soma, but not in the axons, limits the amount of T3 that is transferred from the endosomal/NDL vesicles to the cell nucleus.

### T3 Transport Triggers Localized TH Signaling in the Mouse Brain

Next, we looked for *in vivo* evidence supporting the idea that axonal transport of T3 can trigger TH signaling. We did this by revisiting our previous hypothesis that retrograde axonal transport of T3—–from the median eminence to the hypothalamic paraventricular nucleus (PVN)—–explains how T3 generated by tanycytic D2 can down-regulate TRH expression (*39*). This was experimentally tested here by injecting T3^I125^ directly into the hypothalamic median eminence (ME) of rats. Thirty minutes later, radioactivity was detected in the PVN, whereas only background was found in the lateral hypothalamus and cerebral cortex (**Fig. 6A-F**). Next, we test whether axonal T3 transport plays a role in TH signaling the TH-action indicator THAI transgenic mouse (*27*); in this mouse, the cerebral cortex and hypothalamus are highly responsive to T3 (**Fig. 6D**; **Fig. S4A**). To test for axonal T3 transport and action, we stereotaxically implanted T3 crystals into the primary motor cortex (M1) of the right hemisphere (**Fig. 6D**), and after 48 h we not only found local induction of Luc mRNA (2.7 ± 1.2-fold) at the site of implantation but also induction in the contralateral M1, which receives interhemispheric axonal projections through the corpus callosum (**Fig. 6E**). In addition, we also detected T3 signaling in the ipsilateral secondary somatosensorial cortex (S2) that likewise receives projections from M1 (*42*). Of note, the absence of induction of two T3-responsive markers in the hypothalamus, Luc (**Fig. S4C**), and the TRH-degradation enzyme (trh-de; **Fig. S4C**), a highly T3-sensitive region not connected directly with M1, indicates that the implanted T3 molecules did not diffuse randomly. These findings support the hypothesis that T3 can be selectively transported along neuronal axons and can, therefore, initiate TH signaling in distant but discrete brain areas.

**Fig. 6.**
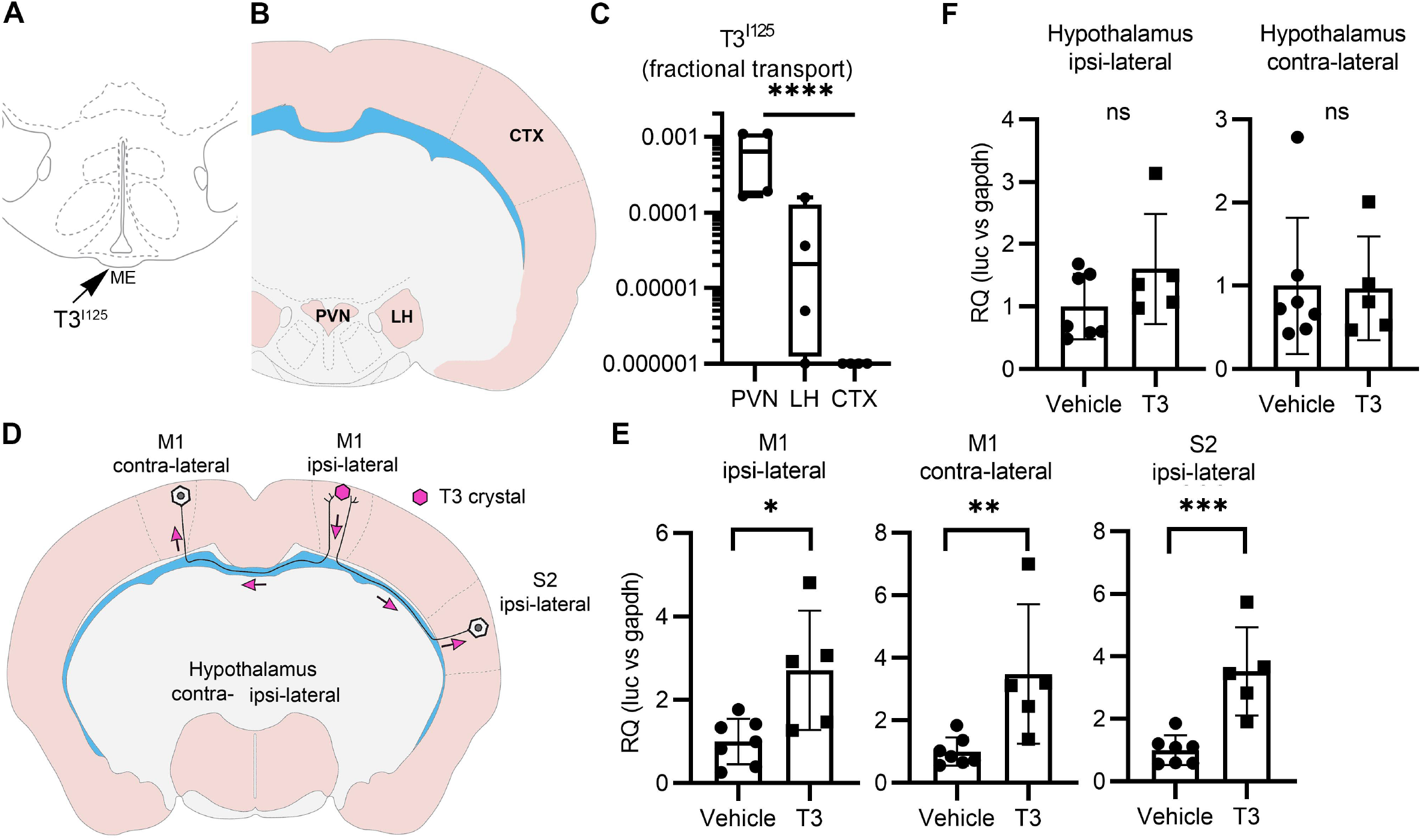
Axonal T3 transport can initiate TH signaling between two interconnected brain areas. **A.** Cartoon showing the site of injection of T3^I125^ in the median eminence (Bregma – 2.56mm). **B.** The areas dissected after 30 minutes of injection are indicated (Bregma −1.80). **C.** Quantitation of the transported T3^I125^ from the median eminence to the indicated areas. **D.** A T3 crystal (pink hexagon) was inserted into the M1 of the right hemisphere, which receives axons from neurons located in the indicated areas. The pink arrows indicated the direction of the transported T3. **E.** Quantitation of Luc mRNA levels at the indicated brain areas. **F.** Hypothalamic expression of Luc mRNA. Values are mean ± SD of 4-7 independent experiments; \**P* <0.05, \*\**P* < 0.01, \*\*\**P* < 0.001; ns: non-significant.

## Discussion

The discovery that astrocyte-derived T3 acts in a paracrine fashion to regulate gene transcription in neighboring neurons placed DIO2 at the crossroads of therapy for hypothyroidism (*43*). Here we show that a common DIO2 polymorphism (Thr92Ala-Dio2) halves astrocytic T3 production, leading to a localized reduction in TH signaling in the hippocampus, which is likely to explain the cognitive phenotype exhibited by mice carriers of the Ala92-Dio2 allele (*30*). Therapy containing L-T3 is one of the alternatives to improve the effectiveness of TH replacement therapy, but critical questions remain unanswered (*44*). Here we addressed some of those questions and provide the mechanistic basis for the T3 signaling in the brain including the roles played by MCT8 and D3. We now show that neurons utilize the coordinated expression of MCT8 and D3 to create an unimpeded pathway for T3 to reach the cell nucleus that starts at the axonal termini, where T3 molecules are concentrated into endosomes/NDL. These T3-loaded vesicles are retrogradely transported to the neuronal cell body, delivering T3 to the nucleus where it regulates gene expression. T3 molecules that escape the endosomes/NDL prematurely or enter the cytosol directly from the extracellular space are rapidly inactivated via D3.

Three steps characterize the retrograde transport of T3: First, T3 is loaded into clathrin-dependent MCT8- and D3-containing endosomes/NDL in long distal axons; once inside the endosomes/NDL, T3 is protected from D3-mediated catabolism because D3’s active center is cytosolic (*39*). This is illustrated by the fact that there is no T3 metabolism in MC-AS and adding XH to the MC-AS did not affect D3-mediated T3 catabolism (**Fig. 5C,D**). Second, the T3-containing endosomes/NDL travel retrogradely through a microtubule-dependent mechanism, which is illustrated by the fact that adding CO to the MC-AS decreases by >50% the amount of T3 found in the MC-CS (**Fig. 4C**; **Fig. S2C**). Third, T3 exits the endosomes/NDL and reaches the nuclear compartment to establish a transcriptional footprint. Indeed, we were able to identify retrograde transport of T3^I125^ to the neuronal nuclei (**Fig. 4E**). T3 molecules that bypass this pathway and enter the cytosol directly through MCT8 are subject to active D3-mediated inactivation to T2 (**Fig. 5E**).

In neurons, signaling endosomes/NDL are normally organized at the axonal termini, adjacent to the post-synaptic membrane or glial cells. These endosomes/NDL are retrogradely transported via microtubule-dependent dynein motors from the distal end of a long axon to the cell body, enabling extracellular molecules to modify gene expression (*45*). Endosomes/NDL direct molecular cargo along four main routes: recycling to the cell surface, transport to the Golgi apparatus, degradation in endolysosomes, or transport to the nucleoplasm. The presence of small amounts of MCT8 (**Fig. 3E,F**) and D3 (*18*, *33*) in the nuclear membrane suggests that the latter mechanism is involved in the transfer of T3 to the cell nucleus, allowing for extracellular signals to affect nuclear events such as gene expression (*46*). Our data indicate that not only T3 is cargo to these endosomes/NDL but also that T3 uses this transport system to regulate gene expression (**Fig. 4F**).

The widespread MCT8 expression in the brain (*8*, *47*) supports a role for MCT8 in neuronal T3 signaling, but the exact mechanism remained elusive. Studies using iPSC-derived neural cells suggest that MCT8 might have a greater role at the BBB, a view that is also supported by studies in zebrafish (*47–49*). However, given the high D3 activity in neural cells (**Fig. 5G**), it is challenging to study T3 cellular uptake without considering T3 metabolites. In addition, there are discrepancies between human and animal models because of the expression of alternative TH transporters in the latter (*13*). Thus, a unifying hypothesis for the contribution of MCT8 to TH signaling in neural cells and how its loss-of-function mutations relate to the patients’ neurological manifestations was missing. The present studies identified two important, unequivocal roles played by MCT8 in murine cortical neurons. First, MCT8 is critical in the endosomes/NDL pathway that retrogradely transports T3, including the exit of T3 from the vesicles (**Fig. 4F-H**,), which can then enter the cell nucleus and affect gene transcription. Second, as in other cells, MCT8 mediates T3 transport into the cytosol (cell bodies; **Fig. 5G**).

The brain of a healthy, non-pregnant adult exhibits the highest D3 activity level (*50*). Nonetheless, >90% of the TRs in the brain are occupied with T3 (*7*), a figure much higher than any other tissue — TR occupancy with T3 in the liver is about 50%. Thus, the present findings resolved this paradox, explaining how a pathway that takes advantage of the topological orientation of D3 (catalytic active center facing the cytosol) avoids catabolism of T3 that is incoming through the endosomes/NDL. The physiologic implications of these findings are considerable, as they reveal potential checkpoints for TH signaling in neurons. For instance, we had previously observed that in hypoxic neurons D3 accumulates in the nuclear membrane, reducing TH signaling (*33*). Accordingly, we now show that inhibiting D3 in the cell body (nucleus) enhances the transfer of T3 from the endosomes/NDL to the cell nucleus and stimulates gene expression (**Fig. 4F**). Thus, under certain pathological conditions (e.g. hypoxia) the entry of T3 in the nucleus can be affected by the presence of D3.

The present studies also advance our understanding of how T3 in the median eminence (plasma-born and locally D2-generated) may regulate TRH expression in the PVN, expanding on our original hypothesis (*39*). The speed with which injected T3 travelled retrogradely to the PVN—just 30 min—is remarkable. In addition, the present studies brought to light the unanticipated reality that T3 molecules can be taken up by long neurons and transported to distant locations in the brain. In fact, we found that T3 originating from one brain hemisphere was taken up, transported, and had an effect on gene expression in neurons located in the contralateral hemisphere (**Fig. 6,D,E**). Of note, the intensity of the T3-induced Luc expression varied between ~2.7-3.4-fold (M1 ipsilateral and contralateral; **Fig. 5E**), which is within the fold-range obtained during the transition between hypothyroidism and TR saturation in the cerebral cortex (2.5-fold; **Fig. S4B**). This suggests that the axonal pathway that retrogradely transports T3 operates well within the physiological context.

The present study can be expanded in the future to address some important remaining gaps. The mechanism through which T3 is taken up and concentrated within the endosomes/NDL has not been established. Despite clear evidence of MCT8 in the nuclear membrane, details of the transfer of T3 from the endosomes/NDL to the nucleus need to be clarified. Lastly, the possibility that other TH transporters, e.g. LAT1-2, play similar roles as MCT8 needs to be investigated. This is less likely to be relevant given that SC fully prevented the T3 (MC-AS→MC-CS) induction of Luc (**Fig. 4F**).

The present findings revealed that carrying the Ala92-Dio2 polymorphism reduces the effectiveness of therapy with L-T4 in mice with hypothyroidism because of decreased T3 production in the hippocampus. The therapeutic effect of treatment with L-T3 is explained because T3 molecules that are selectively taken up into clathrin-dependent, MCT8- and D3-containing endosomal/NDL, are protected against degradation during the transport to the cell nucleus. This pathway contrasts with T3 entering the cytosol of neurons directly from the cellular compartment, which can be rapidly inactivated to T2. In addition, perikaryonal D3 modulates the entry of T3 molecules transitioning from the endosomes/NDL and those entering the cytosol directly from the extracellular space. Altogether, the present findings resolve the paradox of the high T3 nuclear content in the brain amid a very high level of D3 activity. They explain how therapy for hypothyroidism that contains L-T3 can bypass DIO2 defects in astrocytes and safely reach the neuronal nucleus to restore TH signaling.

## Methods

All experiments were approved by the Institutional Animal Care and Use Committee at the University of Chicago (#72577) or by the Animal Welfare Committee at the Institute of Experimental Medicine and followed the American Thyroid Association Guide to investigating TH economy and action in rodents and cell models (*51*).

### Animals

THAI mice (*27*), homozygous THAI-Thr92- and THAI-Ala92-Dio2 mice, and Wistar rats (Charles River, Hungary) were housed in temperature-controlled (21 ±1°C) facilities, with automated light and dark cycles of 12 h, fed ad libitum with chow diet and studied at the indicated ages. Unless otherwise specified, male THAI-Thr92- and Ala92-Dio2 mice, 8-to 10-week-old animals were used throughout experiments. Hypothyroidism was induced by adding MMI (0.05%) to the drinking water and a low iodine diet for 7 weeks (*30*, *52*). Some animals received an L-T4 treatment given at 1.7 and 1.85 ug/100g BW/day via gavage, for 2 weeks.

### Primary cultures of neonatal astrocytes from the cortex, hippocampus, and cerebellum

Primary astrocyte cultures (*53*) were established from the whole neocortex, hippocampus, and cerebellum of post-natal day 3 (P3) Thr92- and Ala92-Dio2 mice. The pups were decapitated and the brain and cerebellum dissected. The meninges and blood vessels were removed, and the cerebral cortices and hippocampi were isolated and incubated for 30 min in DMEM with 0.25 % tryp-sin-0.53 mM EDTA. To obtain a single-cell suspension, tissues were dissociated by triturating with a 10 ml followed by a 5 ml serological pipette and then passed through a 50 μm cell strainer. After centrifugation at 1,200 rpm (5 min), the pellet was suspended in astrocyte-culture medium (ACM; DMEM high glucose containing 10% FBS and 1% penicillin-streptomycin). The cells obtained from each region were seeded at low density in 2 wells of collagen-coated 6-well plates (150,000 cells per well). Cells were incubated for 7 days with medium changes every 72 hours. Once cultures reached 80% confluency, cells were split following a 1:3 ratio, so that each region was cultured in 6 wells. Experiments were done when the cells after the first passage reach a confluency of 80%.

### T3^I125^ injection in the median eminence

220 g adult Wistar male rats were kept on a heating pad while undergoing ventral transsphenoidal surgery (*54*) to expose the median eminence (ME) of the hypothalamus. The T3^I125^ was freshly purified by LH-20 column and applied to the ME using a Nanoliter 2010 microinjector. ~40,000 cpm were injected in a volume of 50 nl. After 30 min, the animals were killed, the brain was removed and PVN, the lateral hypothalamus (LH), and a cortical sample were microdissected with the Palkovits’s punch technique (*55*) and counted in a gamma counter.

### T3 signaling in the THAI mouse brain

2-month-old male THAI mice were injected with 0.1 and 1 μg/BW T3 i.p. and decapitated 24 h later. The cerebral cortex was dissected and assayed for Luc activity as described (*27*) using an assay system reagent (Promega, Madison, WI) on a Luminoskan Ascent luminometer (Thermo Electron Corp. Labsystems, Vantaa, Finland); the relative light unit (RLU) was normalized to protein content. To study T3 trafficking across different cortical areas, we inserted crystalized T3 (*56*) into the M1 of one hemisphere using stereotaxic surgery; control mice were sham-operated. Animals were decapitated after 48 h. The M1 and the contralateral M1, the ipsilateral S2, and the ipsi- and the contralateral portion of the hypothalami were microdissected and processed for RT-qPCR using TaqMan Real-Time.

### Primary embryonic cortical neurons were cultured in a microfluidic compartmentalized device

PCNs were isolated from E16.5 THAI mouse embryos. Briefly, embryos were removed, and the cerebral cortex dissected, stripped of meninges, and dissociated into a combination of Ca^2+^ and Mg^2+^ free Hanks balanced salt solution (HBSS) containing 0.25 % trypsin-0.53 mM EDTA, then mechanically triturated using fire-polished glass Pasteur pipettes. Isolated cells were passed through a 40 μM cell strainer to reach a cell density of 4.5 × 10^6^ cells/ml. We used a microfluidic compartmentalized culture device that contains 450 μM long microchannels connecting MC-CS and MC-AS and permits only distal axons to grow into the MC-AS (XonaChip, Cat# XC450, Xona Microfluidics Temecula, CA, USA). The cortical neurons were plated at a density of 5-9 x 10^4^ cells/device in the cell compartment with a growth medium composed of the neurobasal medium, 2 % B-27 supplement, 1 % GlutaMax, and 1 % antibiotic-antimycotic (Penicillin-streptomycin; all from Gibco). On DIV ~ 2, one-half of the medium was replaced with a growth medium containing the anti-mitotic cytosine arabinoside (Sigma-Aldrich) which restricts astrocytes and microglia to <0.01% (*57*). Thereafter, the growth medium was replaced every other day. During the different experiments, each compartment was fluidically isolated by hydrostatic pressure, accomplished by keeping the medium volume in one compartment higher than in the opposite compartment, allowing us to differentially treat either side(*58*). For the isolation of DRG neurons, the dorsal root ganglia were dissected and cultured from 2-day-old Wistar rats and THAI 6-day-old THAI mice according to published protocols(*59*, *60*)All other procedures were as with the PCNs.

The compartments remained fluidically isolated (Suppl. Fig. 2D). In the absence of cells, adding T3^I125^ to the MC-AS revealed the predominant T3^I125^ peak after 24-72h (minimal T2^I125^ and I-^I125^ peaks were also detected) in the MC-AS chromatograms (Suppl. Fig. 2E, orange panel), whereas only background radioactivity was detected in the MC-CS (Suppl. Fig. 2E, blue panel). Even in the presence of cells, no signal was detected on the MC-CS after 24 h of adding the fluorescent dye (Alexa Fluor 594 hydrazide) in the MC-AS (Suppl. Fig. 2C).

### Cell staining and Immunofluorescence studies

Cultures at DIV ~ 10 were fixed in 4% paraformaldehyde for 20 min, washed twice in PBS, and then permeabilized in PBS with 0.1 % Triton X-100 for 5 min. Cultures were blocked in PBS with 5 % BSA for 15 – 30 min at room temperature and incubated with primary antibody diluted in PBS, at 4°C overnight. Cultures were rinsed 3 times and incubated for 2 h at room temperature with a secondary antibody. Primary and secondary antibody dilutions were as follows: mouse monoclonal anti-Lamp1 antibody 1:1000 (Sigma), rabbit polyclonal anti-MCT8 antibody 1:400 (Atlas antibodies), rabbit polyclonal anti-D3 antibody (Novus Biologicals), rabbit polyclonal anti-GFAP (1:250), alexa 488 conjugated goat anti-mouse IgG 1:200 (Vector), alexa 594 conjugated horse anti-rabbit IgG 1:200 (Vector), alexa 488 conjugated horse anti-rabbit IgG 1:200 (Vector). The images were analyzed by NIS-Element AR (Nikon Instruments) or ImageJ software (NIH). Final Fig.s were prepared on Adobe Photoshop.

### Immuno-electron microscopy for Mct8

Mice were anesthetized with a mixture of ketamine and xylazine (50 and 10 mg/kg BW i.p., respectively), perfused (trans-cardiac) with 10 ml 0.01 M phosphate-buffered saline (PBS), and fixed with 40 ml of 2% paraformaldehyde and 4% acrolein in 0.1 M phosphate buffer (PB). The brains were removed and postfixed by immersion in 4% PFA in PBS overnight at room temperature. Coronal 25-μm thick sections containing the primary somatosensorial cortex were cut with a vibratome (Leica VT 1000S) and stored at −20°C in 30% ethylene glycol, 25% glycerol in 0.05M PB until further use. Pretreatment included 30 min incubation with 1% sodium borohydride and 15 min with 0.5 % H_2_O_2_, followed by a sucrose gradient (15 → 30 %) and three frozen-thaw cycles in liquid nitrogen. For immunohistochemistry, sections were blocked with 2% normal horse serum and incubated with rabbit polyclonal antiserum against MCT8 (1:20,000; kind gift of Dr. TJ Visser) for 4 days at 4 °C, followed by biotinylated donkey anti-rabbit IgG (1:200; Jackson Immuno Research Labs,) for two hours and 0.05 % DAB / 0.15 % Ni-ammonium-sulfate / 0.005 % H_2_O_2_ in 0.05 M Tris buffer (pH 7.6). The staining was silver-gold-intensified using the Gallyas method(*41*, *61*). For electron microscopy, sections were incubated in 1% osmium-tetroxide for 1 h at room temperature and then treated with 2% uranyl acetate in 70% ethanol for 30 min. Following dehydration (ethanol - acetonitrile) the sections were embedded in Durcupan ACM epoxy resin on liquid release agent coated slides and polymerized at 56 °C for 2 days. Ultrathin, 60 – 70 nm-thick sections were cut with Leica UCT ultramicrotome (Leica Microsystems, Vienna, Austria), were mounted on Formvar coated, single-slot grids, and treated with lead citrate. Images were obtained using a transmission electron microscope (JEOL-100 C).

### Lysosomes isolation by ultracentrifugation

We used a lysosome enrichment kit (Thermo Scientific) and followed the manufacturer’s instructions. Briefly, approximately 50-200 mg of cells were harvested and lysed using a sonicator (15 bursts; 9W power). Subsequently, the homogenate was centrifuged to remove cellular debris. The resulting supernatant was overlayed on several discontinuous gradients of the OptiPrep Cell Separation Media (60% iodixanol in water with a density of 1.32 g/ml) and ultracentrifuged (145,000 *g* for 180 min at 4° C) to isolate and enrich for lysosomes. The different fractions were removed from the gradient, pellet by centrifugation, and washed three times with PBS. The final pellet was lysed in 30 ul of 0.02M ammonium acetate + 4% methanol + 4% PE buffer and studied through UPLC.

### Iodothyronine chromatography using UPLC

PCNs at DIV ~10 were incubated with 10^6^ cpm of T3^I125^/ml, totaling 60 ul per well (total two wells). After 24 and 72 h, 100 ul of the medium was sampled, mixed with 100 ul of 0.02 M ammonium acetate + 4% methanol + 4% PE buffer (0.1M PBS, 1mM EDTA), and applied to the UPLC column (AcQuity UPLC System, Waters). Fractions were automatically processed through a Flow Scintillation Analyzer Radiomatic 610TR (PerkinElmer) for radiometry. The D3-mediated deiodination was calculated by the production of T2 ^I125^/ h / mg protein. Primary astrocyte cultures at 80% confluency were incubated with 10^6^ cpm of T4^125^/ml, totaling 500 ul per well. After 24, the medium was collected and the D2-mediated deiodination was calculated by the production of T3^I125^/ h / mg protein.

### TaqMan Real-Time quantitative PCR

Total RNA was isolated from microdissected specific brain areas with NucleoSpin RNA kit (Macherey-Nagel)) or from neurons growing in MC-CS with an RNeasy Mini kit (ThermoFisher). DNA contaminants were digested with DNASE I (Ambion). Undiluted total RNA (1 ug) was reverse transcribed with the High-Capacity cDNA Reverse Transcription Kit (ThermoFisher Scientific, Waltham, MA. cDNA concentration was determined with Qubit ssDNA assay kit and 10 ng cDNA was used in each Taqman reaction. Luciferase expression was detected using a specific TaqMan probe (Applied Biosystems; Assay ID: AIY9ZTZ) using TaqMan Fast Universal PCR Mastermix (Applied Biosystems) and compared with hypoxanthine phosphoribosyltransferase 1 (Hprt1) or Glyceraldehyde 3-phosphate Dehydrogenase (Gapdh) housekeeping genes expression. Reactions were assayed on a Real-Time PCR instrument (Applied Biosystems, Waltham, MA). For *trh-de* gene, we used the #Mm00455443_m1 Taqman probe.

### Statistics

All data were analyzed using Prism software (GraphPad). Unless otherwise indicated, data are presented as scatter plots depicting the mean ± SD. Comparisons were performed by a 2-tailed Student’s t-test, and multiple comparisons were by ANOVA followed by Tukey’s test. A p<0.05 was used to reject the null hypothesis.

## Author contributions

FS-L conducted *in vitro* experiments with mouse cortical neurons and astrocytes, prepared figures, and analyzed data. BB, TF and FS-L conducted *in vivo* experiments with L-T4 treated THAI mouse. RS, PE, KR and YR, CSF and BG conducted *in vivo* experiments and *in vitro* experiments with dorsal root ganglia neurons. FS-L and CF interpreted the data and prepared the manuscript. BG and ACB interpreted the data, prepared the manuscript, and directed all the studies.

## Acknowledgments

The authors are grateful to support from NIDDK DK58538, DK65055, the Hungarian National Brain Research Program 2, NRDIO K125247; FS-L was supported in part by grant DK15070. The technical help of Andrea Juhász and Dóra Fazekas is gratefully acknowledged. ACB is a consultant for AbbVie, Synthonics, Sention, and Thyron.

---

**Fig. S1**. **A system to study TH signaling in neurons. A.** Microfluidic device showing the MC-CS in blue and the MC-AS in orange. **B**. Typical neuronal growth at DIV ^~^ 10, the MC-AS is densely populated by axons **C**. Applying Alexa Fluor 594 in the MC-AS demonstrate fluidic isolation. Inset shows calcein fluorescent axons. **D, E**. In the absence of cells, T3^I125^ is applied in the MC-AS, and only background radioactivity was detected in the MC-CS. Chromatograms from the MC-AS (orange) show typical peaks of T3^I125^ T2^I125,^ and I^I125^.

**Fig. S2. T3 trafficking in rat DRG cells in microfluid chambers. A.** ^~^100.000 cpm freshly purified T3^I125^ was added into the MC-AS. **B.** Rat DRG cells in microfluid chamber transfected with GFP. **C.** Media in the MC-CS was counted on a gamma-meter at the indicated time points and conditions. **D.**Western blot on cultured rat DRG cells using an anti-MCT8. **D** Presence of D3 was confirmed by PCR.

**Fig. S3. Adding SC or XH in the MC-AS had no effect on the MC-CS. A.** T3^I125^ was applied in the MC-CS. **B**. D3-mediated T2^I125^ production in the indicated conditions. Values are mean ± SD of 3 – 6 independent experiments; \*\**P* < 0.01 in comparison with T3^I125^ incubation.

**Fig. S4. T3 triggers TH signaling in the brain of the THAI mice A.** Luciferase mRNA levels in the mediobasal hypothalamus (MBH) and cortex **B.** Quantitation of the Luciferase activity in the cerebral cortex of the THAI mice at the indicated conditions. **C.** Hypothalamic expression of Luc mRNA and Trh-de. Values are mean ± SD of 4-7 independent experiments

