## Supplementary figures and images for "A PATHWAY FOR T3 SIGNALING IN THE BRAIN TO IMPROVE THE VARIABLE EFFECTIVENESS OF THERAPY WITH L-T4"

### Supplemental Figure 1

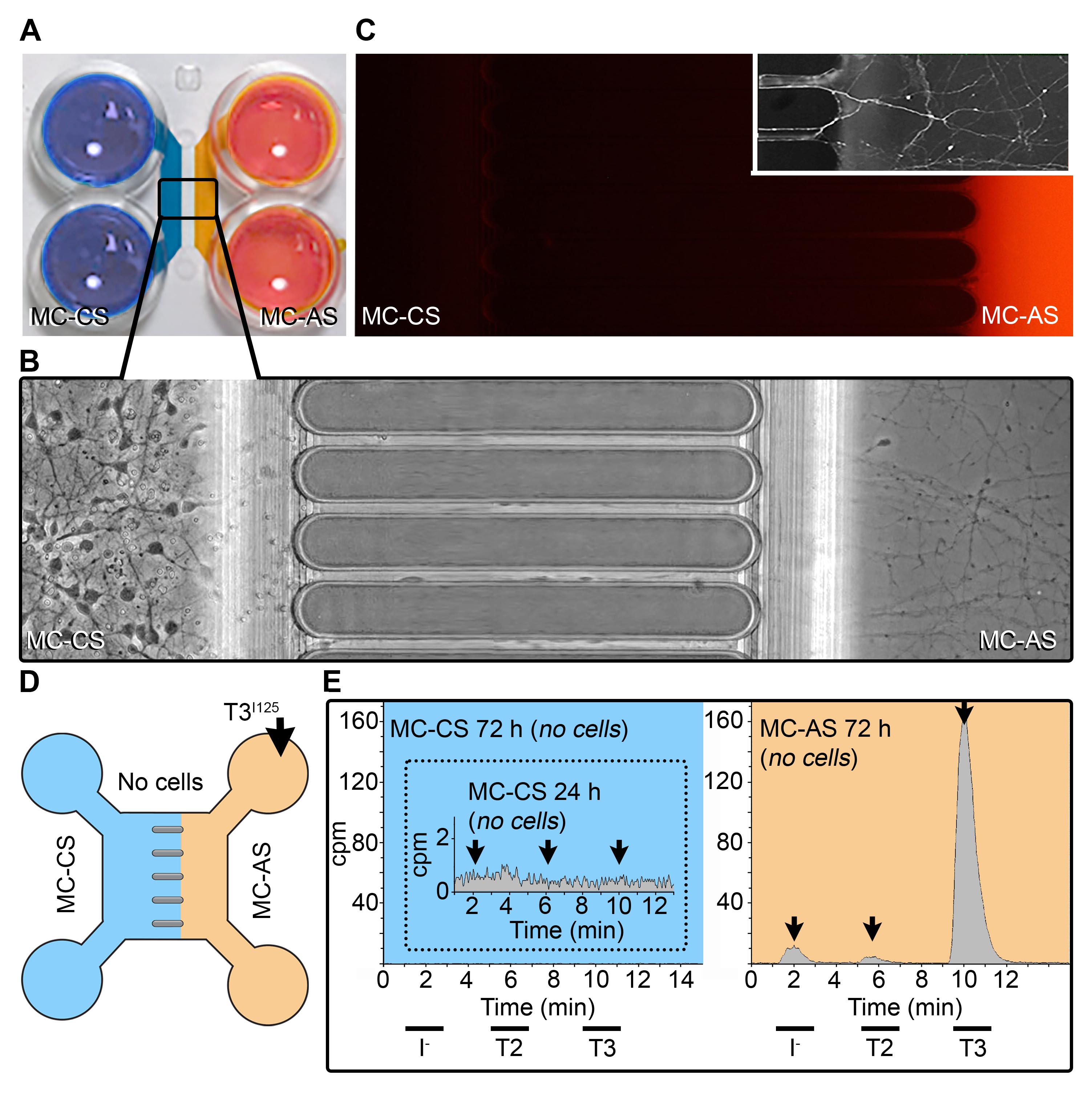

### Supplemental Figure 2

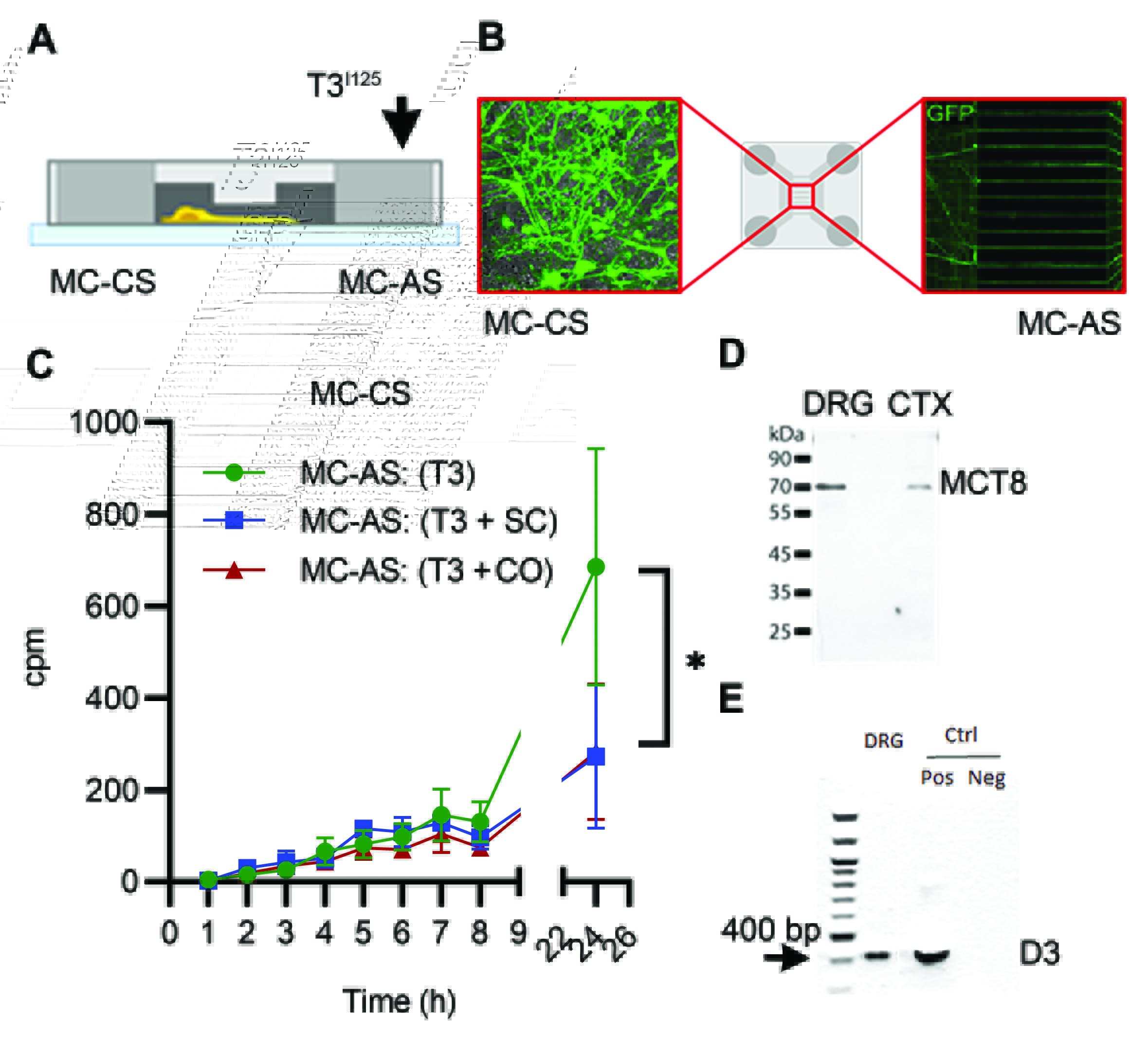

### Supplemental Figure 3

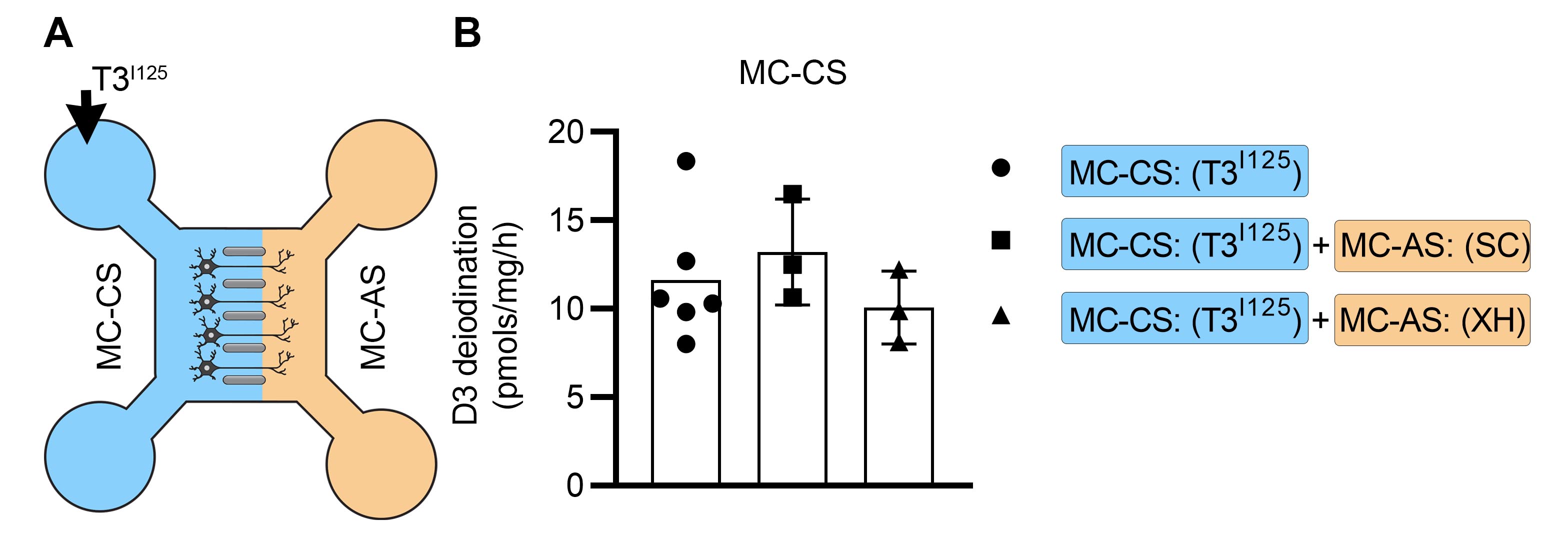

### Supplemental Figure 4

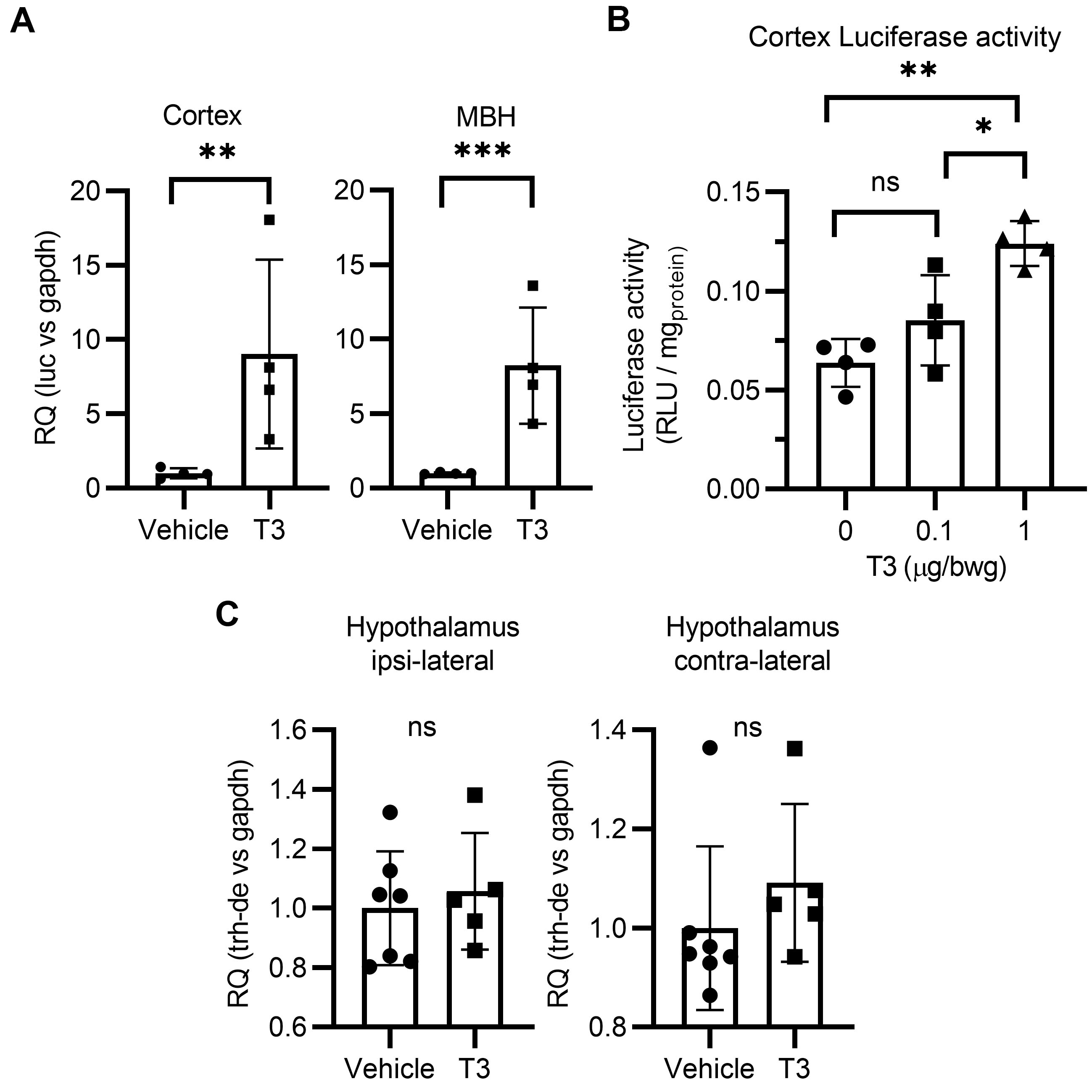
